# Insights into the evolution of herbivory from a leaf-mining, drosophilid fly

**DOI:** 10.1101/2022.12.07.519390

**Authors:** Jessica M. Aguilar, Andrew D. Gloss, Hiromu C. Suzuki, Kirsten I. Verster, Malvika Singhal, Jordan Hoff, Robert Grebenok, Paul D. Nabity, Spencer T. Behmer, Noah K. Whiteman

## Abstract

Herbivorous insects and their host plants comprise most known species on Earth. Illuminating how herbivory repeatedly evolved in insects from non-herbivorous lineages is critical to understanding how this biodiversity is created and maintained. We characterized the trophic niche of *Scaptomyza flava*, a representative of a lineage nested within the *Drosophila* that transitioned to herbivory ∼15 million years ago. We used natural history studies to determine if *S. flava* is a true herbivore or a cryptic microbe-feeder. Specifically, we quantified oviposition substrate choice and larval viability across food-types, trophic-related morphological traits, and nitrogen isotope and sterol profiles across putatively herbivorous and non-herbivorous drosophilids. We confirmed that *S. flava* is an obligate herbivore of living plants. Paired with its genetic model host, *Arabidopsis thaliana, S. flava* is a novel and powerful system for exploring mechanisms underlying the evolution of herbivory, a complex trait that enabled the exceptional diversification of insects.

## Introduction

Identifying the macro- and micro-evolutionary factors that create and maintain biodiversity is a central goal of evolutionary biology. Studies elucidating the ecological interactions between plants and their insect herbivores, the two most species-rich groups of macroscopic organisms on the planet, have been critical to addressing this challenge. It is well established that interactions between plant host and insect herbivores drive increased diversification rates in insects (Ehrlich and Raven 1964; Mitter, Farrell, and Wiegmann 1988). Yet, there are also challenges to herbivory as a life history strategy: although 50% of insects are herbivorous, only 30% of extant insect orders contain herbivores (Schoonhoven, van Loon, and Dicke 2005; Mitter, Farrell, and Wiegmann 1988). This paradox—that a minority of orders contain nearly half of all insect species—suggests that plants present challenging evolutionary hurdles for insects to initially overcome as food and habitat (Southwood 1972).

There has been progress in understanding the genetic basis of herbivory, speciation, and host-plant specialization in herbivorous insects (Feder, Chilcote, and Bush 1988; McPheron, Smith, and Berlocher 1988; Gompert et al. 2012; Heliconius Genome Consortium 2012; Soria-Carrasco et al. 2014; Goldman-Huertas et al. 2015; Vertacnik and Linnen 2017). However, because herbivory evolved over 50 million years ago in the most diverse herbivorous insect lineages, the development of new model systems in which herbivory has evolved more recently is critical to understand broad patterns underlying the evolution of herbivory itself.

An ideal lineage in which to study the evolution of herbivory is one in which: (1) this trait has evolved relatively recently (so that close, non-herbivorous relatives can be used in comparative studies), (2) genomic tools are available (a well-annotated reference genome; amenability to germline transformation), (3) its generation time is relatively short (to facilitate laboratory and field studies), and (4) it attacks a rapidly cycling host plant with its own suite of well-developed tools (so that both sides of the plant-herbivore equation can be experimentally manipulated). The drosophilid fly *Scaptomyza flava* (Fallén, 1823) meets all of these criteria (Whiteman et al. 2011, 2012; Goldman-Huertas et al. 2015; Gloss et al. 2014; Matsunaga et al. 2022; Peláez et al. 2022).

The transition from microbe feeding to herbivory by the lineage containing *S. flava* is relatively recent (∼15 million years before present) among an array of insect species with sequenced genomes (Supplementary Figure 1). It is nested in a clade of well-studied, non-herbivorous drosophilid flies, including *Drosophila melanogaster*, and *S. flava* attacks the model plant *Arabidopsis thaliana* in nature and the laboratory (Whiteman et al. 2011; Mitchell-Olds 2001). The adult females have leaf-puncturing ovipositors that they use to create feeding punctures and lay eggs in leaves, where larvae hatch and complete development (Peláez et al. 2022). The constitutive and jasmonate-induced defenses of *A. thaliana* provide some basal and inducible resistance against *S. flava* (Whiteman et al. 2012). However, the trophic niche of *S. flava*, a key aspect of its natural history, has not yet been empirically characterized.

*S. flava* and congeners are closely associated with living vegetative tissue of plants (N. A. Martin 2004; Hackman 1959), and can only be reared from living plants (Whiteman et al. 2011), but these factors alone do not eliminate the possibility that *S. flava* and its relatives are cryptic microbe feeders and thereby gain nutrition from this trophic level. The possibility that *S. flava* is gaining most of its nutrition from microbes on and within leaves is especially relevant since *Scaptomyza* is nested within an ancestral microbe feeding lineage in which most species consume microbes and associated decaying plant tissue. In addition, the gut microbial community in *S. flava* overlaps almost completely with the phyllosphere bacterial community of its host plant (O’Connor et al. 2014). This overlap in community composition is consistent both with an animal acquiring nutrients sufficient for development from these microbes and with an herbivore that may ingest microbes incidentally.

### What makes an herbivore?

Herbivory is a complex organismal syndrome that involves a suite of inter-dependent adaptations. In the most fundamental sense, an herbivore is an organism that obtains most or all of its nutrients from living plant tissues. There are several subsidiary traits that make herbivory possible; these traits evolve from ancestral character states already shaped by acquiring nutrients from other sources.

For most immature herbivorous insects, initial access to food largely depends on maternal oviposition choices. These choices are informed by visual (Lyu et al. 2018; Nagaya, Stewart, and Kinoshita 2021) and chemosensory (Goldman-Huertas et al. 2015; Matsunaga et al. 2022) cues relaying information about environmental conditions. Although ovipositing females generally lay eggs where larval survival is expected to be high, there are some instances where eggs are laid on less suitable hosts, for example, when more suitable hosts are rare, or the costs to egg laying are low (i.e. when a female has high fecundity) (Jaenike 1978). In these cases, oviposition behavior is one important factor that drives host shifts (Bernays and Chapman 1994).

Additionally, given the differences in the physical properties of food substrate and nutrient availability in plants, morphological adaptations to plant feeding are also common. Herbivores tend to have longer gut lengths compared to non-herbivorous close relatives, which may increase digestion efficiency. This pattern has been observed in many different animal lineages, including crabs, grasshoppers, lizards, and fish (Griffen and Mosblack 2011; Yang and

Joern 1994; Vervust et al. 2010; German and Horn 2006). Another commonly observed morphological adaptation that facilitates trophic niche shifts are changes in the morphology of the mouthparts. In drosophilids, plastic changes in maxillary mouth hook dentation accompany shifts from microbe-eating to cannibalism (Vijendravarma, Narasimha, and Kawecki 2013), and in lizards, changes in dentition have also coincided with shifts from carnivory to herbivory (Vervust et al. 2010).

Although behavioral and morphological adaptations are associated with trophic niche shifts, herbivores must survive primarily on nutrients supplied by their host plants. All organisms require a common set of essential nutrients, but microbes, plants, and animals differ in the ways these nutrients are integrated depending on their trophic niche. These differences in nutrient integration translate to differences in nutrient composition in organisms across trophic levels.

Two salient nutrients in this regard are sterols and protein (Behmer and David Nes 2003; Jing and Behmer 2020). Sterols are lipids that are essential for insect development, but because insects cannot synthesize them *de novo*, they rely on food-sources to obtain them or supplement their diet (Clark and Block 1959; Li and Jing 2020; Ikekawa, Morisaki, and Fujimoto 1993).

Insect sterol profiles are shaped both by initial sterol profiles of food sources, as well as variation in the types of sterols that can be synthesized through dealkylation (Janson et al. 2009; Jing, Grebenok, and Behmer 2013; Jing et al. 2012). Similarly, protein (amino acids) is an essential and often limiting nutrient, and ratios of heavy to light nitrogen isotopes (δ^15^N) from protein in food can be used as indicators of trophic level (Perkins et al. 2014). As fixed nitrogen moves through trophic levels, light isotopes (14N) are preferentially lost resulting in an enrichment of heavier isotopes (15N) in animals in higher trophic levels. When comparing insects that eat microbes to insects that eat plants, a relative enrichment in heavy nitrogen isotopes occurs (McCutchan et al. 2003).

To determine the feeding niche of *S. flava* (Hackman 1959), we compared a subset of trophic niche-related traits between *S. flava* and closely related microbe feeding relatives, including *S. pallida, S. hsui* and *D. melanogaster* (Figure 1). *S. pallida* is in the subgenus *Parascaptomyza* and *S. hsui* is in the subgenus *Hemiscaptomyza* (Lapoint, O’Grady, and Whiteman 2013). Neither of these subgenera of *Scaptomyza* are known to contain herbivorous species, unlike the subgenus *Scaptomyza*, in which *S. flava* nested. We typically rear both *S. pallida* and *S. hsui* in the laboratory from yeast-based Drosophila media with rotting spinach added, suggesting that these two species retain the ancestral character state for the family and eat microbes found in decaying plant tissue. It is well understood that *D. melanogaster* mainly feeds on microbes, especially brewer’s yeast (*Saccharomyces cerevisiae*) and its relatives found in decaying fruit. Salient traits that we compared across this array of four drosophilid species included: 1) oviposition preference and ability to develop on alternative food types representing ancestral dietary substrates–yeast media and decaying plant vegetative tissue, 2) morphological measurements of trophic-related traits, and 3) the metabolic breakdown and integration of food products into the body (sterols and nitrogen isotope ratios). We hypothesized that if *S. flava* is herbivorous, these flies would show a strong oviposition preference for living plants as well as morphological (gut and mouthpart) and nutritive adaptations that differ from non-herbivorous close relatives.

**Figure 1:**
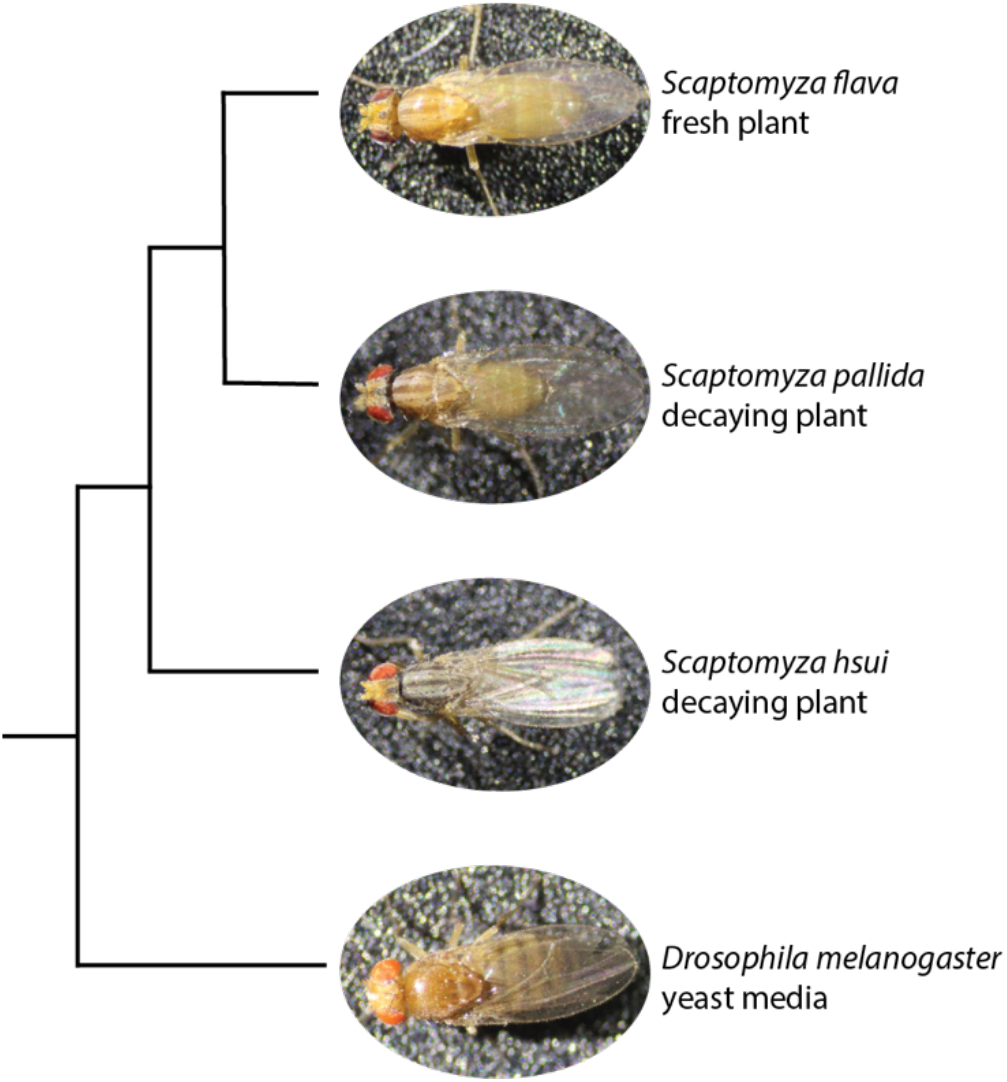
Phylogenetic relationship and laboratory food substrate *of S. flava, S. pallida, S. hsui*, and *D. melanogaster*. Tree topology from Peláez et al., 2022.

## Materials and Methods

### Oviposition choice and development assays

Oviposition preference assays: *S. flava* has exclusively been reared from living plants when collected in nature (Whiteman et al. 2011, 2012; Gloss et al. 2014). To determine if *S. flava* occupies a broader feeding niche than reported previously, we offered adult female flies a choice between living plants, decaying plants, and yeast-seeded media, and compared the choices of *S. flava* to that of microbe-feeding *D. melanogaster*. For both fresh and decaying plant treatments, we grew *A. thaliana* (Col-0) plants to a six-week old rosette-stage as described in Whiteman et al. (2011) in Conviron growth chambers. To facilitate fungal decay that is often associated with *Drosophila* spp. colonization, we froze plants overnight to rupture cell walls, seeded the plant surface with baker’s yeast (Fleischmann’s) and subsequently placed in a warmer environment (approx. 24°C) in the lab for three days before use (Chandler et al. 2012). We prepared standard Drosophila yeast media seeded with the same baker’s yeast in petri dishes that approximated the diameter of the fresh and decaying plant rosette choices. Yeast media was made by the UC Berkeley fly food facility (Supplementary methods). We randomly placed the three food substrates in a 30 × 30 cm mesh cage at 22°C and approximately 60% humidity. We then released four mated, 6-8 day old adult female flies from either an outbred *S. flava* colony collected from wild mustards (*Barbarea vulgaris and Turritis glabra*) near Dover, NH, USA or Canton-S *D. melanogaster* inside the cage. We counted eggs from each substrate after 24 hours. We compared differences in oviposition preference and viability among substrates using a permutation-based Kruskal-Wallis test in the R package *coin* (Hothorn et al. 2008; R Core Team 2021). To ensure inferences were robust to the large number of tied values, we approximated significance against a null distribution constructed by Monte Carlo resampling (N=1e^7^) stratified by experimental block.

*S. flava* viability assays: To determine if *S. flava* requires living plants to complete development, we assayed larval viability on living plants, decaying plants, and yeast-seeded *Drosophila* media, prepared and maintained as described above. Because we found that *S. flava* will only oviposit on living plants, we transferred 3-5 day old larvae from *A. thaliana* to each substrate. We tested nine replicates of six larvae for each substrate, and censused and removed adults that emerged at 13, 20, and 27 days. Differences among substrates were calculated as described above.

#### Comparative morphological measurements

We measured traits that are commonly associated with adaptations to herbivory including gut length, cephaloskeleton size, and mouth hook shape and tooth number in third instar *D. melanogaster, S. hsui, S. pallida*, and *S. flava* larvae. We reared flies on their normal lab diet: *D. melanogaster* on yeast media (UC Berkeley fly facility; Supplementary methods), *S. hsui* on yeast media with decayed spinach (UC Berkeley fly food facility + Earthbound Farms brand spinach; Supplementary methods), and *S flava* on fresh *A. thaliana* plants. We performed all shape and size measurements in imageJ (Schneider, Rasband, and Eliceiri 2012), and for all comparisons, we ran ANOVA and Tukey’s honestly significant difference test (Tukey’s HSD) in R (R Core Team 2021).

We measured wandering third instar larvae gut length and cephaloskeleton size measurements. We estimated this developmental stage by larval size and behavior and confirmed under a dissection scope by examining the anterior spiracles for a branching pattern rather than clubbed (N. A. Martin 2012) and an orange hue on the posterior spiracles. All measurements were done in ImageJ (Schneider, Rasband, and Eliceiri 2012) from photos taken with a scale bar and standardized magnification on a dissecting microscope. We took body measurements when the larvae were in a fully extended state, which we initiated by submerging larvae in 70°C water for ∼5 seconds (Godoy-Herrera et al. 1984). We measured midgut length, as defined from the proventriculus to the Malpighian tubules, by laying the organ such that it was in a single plane, aided by a glass coverslip, while avoiding stretching or compressing. We isolated cephaloskeletons from soft tissue using jeweler’s forceps and took measurements from the tip of the oral process of the mouth hook to the bridge of the posterior process. We divided gut length and cephaloskeleton length by body length to account for variation caused by larvae size.

For mouth hook tooth quantification and shape measurements, we collected the anterior-most section of the third instar mouth hook from the puparial case. We first soaked specimens in 70% ethanol for a minimum of 5 minutes to aid in dissection. We then counted mouth hook tooth number on specimens that we first mounted on glass slides with Fisher Scientific Permount mounting media. We identified the details of the mouth hook teeth using a combination of photographs and visual confirmation under 40x magnification. We mounted right and left sides of the mouth hook separately. Counting and measurements were done on the mouth hook side that was more clearly prepared and/or photographed, or if both sides were of equal quality, one was randomly chosen. We used landmarks as defined in (Wipfler et al. 2013). We took mouth hook height as the distance from the ventral process of the mouth hook to the dorsal process of the mouth hook, and mouth hook length was defined as the distance between the ventral process of the mouth hook to the oral process of the mouth hook. We calculated the ratio of mouth hook height to length to get a sense of the shape distribution of the mouth hook (i.e. whether mouth hooks were long and thin versus more robust in width compared to height). Tooth size was calculated by subtracting the height of the mouth hook measured from the base of the longest tooth to the top of the mouth hook from the height of the mouth hook from the tip of the tooth to top of the mouth hook. To depict shifts in mouth hook morphology across species, we traced representative mouth hook images in Adobe Illustrator and placed in a species tree from (Peláez et al. 2022).

#### Sterol profiling

To better understand differences in sterol utilization from food sources, we extracted and characterized the sterol types and quantities present in *D. melanogaster, S. hsui*, and *S. flava* and the food on which they are reared in the lab. We chose these three species to represent the range of food types of *Scaptomyza* and close relatives. First, we reared flies on their normal lab diet: *D. melanogaster* on yeast media (Bloomington; Supplementary methods), *S. hsui* on yeast media with decayed spinach (UC Berkeley fly food facility + Earthbound Farms brand spinach; Supplementary methods), plain decayed spinach for comparison and *S flava* on fresh *A. thaliana* leaves. We added previously frozen, thawed store-bought spinach to the top of a subset of the yeast media vials. We left all food preparations at room temperature for 7 days, during which time the yeast media with spinach and the plain spinach were able to decay. *A. thaliana* samples were approximately 30 days from planting and were grown as above.

We weighed and homogenized 21 individual *S. hsui* larvae, 13 individual *S. flava* larvae, a pool of 90 *D. melanogaster* puparia (A. M. Martin 2015) and samples of prepared food that had not been eaten. We processed these samples as per Jing et al., 2013 with slight modification. We extracted and processed free and base hydrolysable sterols, and glycosylated sterols (Supplementary methods, (Jing, Grebenok, and Behmer 2013; Feng et al. 2015). We conjugated free sterols and sterols freed following saponification and acid hydrolysis using a BSTFA+TMCS, 99:1 (Sylon BFT) Kit (Supelco, Bellefonte PA). We re-suspended samples in 800 µl of conjugation solution and hexane (4:1 v/v) and incubated for 12 h in the dark at room temperature. Following incubation, we added 2 ml of hexane and 2 ml of 70% methanol:water (v/v) and mixed the sample by pipetting. We removed the hexane fraction, evaporated to dryness under nitrogen. We subjected all fractions to GC-MS analysis.

We plotted the average sterol composition for *S. hsui* and *S. flava*, and the pooled composition for *D. melanogaster* in R (R Core Team 2021)). We calculated the Bray-Curtis dissimilarity between all samples using the vegan package (Oksanen et al. 2022). This measure takes both the presence of sterol compounds and the amounts of sterol compounds into account.

#### Isotope profiling

To measure δ^15^N in *S. flava*, we reared flies in replicate cages with ample *A. thaliana* Col-0 rosettes grown to approximately 6 weeks as described above. We collected plant tissue immediately prior to exposure to adult *S. flava*, and again after 10 days of feeding using leaf tissue adjacent to any mines. We collected 25 third instar larvae for each of three replicates at 10 days and at 21 days. Immediately after collection, we dried tissues at 50**°**C, ground to a fine powder, and combusted using an Elemental Combustion System (Costech Analytical Technologies, Valencia, CA) coupled to a continuous-flow gas-ratio mass spectrometer (Thermo Finnigan Delta PlusXL, San Diego, CA). We quantified δ^15^N using acetanilide standard for elemental concentration and IAEA-N-1 and IAEA-N-2. Precision is better than ± 0.2 for δ^15^N (1 s.d.), based on repeated internal standards.

To compare *S. flava* δ^15^N trophic enrichment with other herbivorous and non-herbivorous organisms, we performed a simple search (in 2013) of “herbivore” “insect”, “trophic”, and “isotope” but only those publications with raw or summary data available on δ^15^N for both consumer and food/host were used. We found additional sources by screening papers that cited those found in the original search. We restricted our search to fluid-feeding and defoliating herbivores. For both literature and lab derived measurements, we calculated δ^15^N trophic enrichment by subtracting diet δ^15^N from insect δ^15^N. We compared δ^15^N trophic enrichment values of herbivores and non-herbivores found in the literature using an ANOVA run in R (R Core Team 2021)).

## Results

### *S. flava* oviposition choice and development assay

In a three-way choice test offering equal proportions of approximately six-week-old, living *A. thaliana* rosettes, decaying *A. thaliana* rosettes, and yeast-seeded nutrient media, *S. flava* laid eggs exclusively in living leaves (N = 34 total eggs across 10 out of 17 trials, chi-squared = 19.767, p = 4.82e^-5^, Kruskal-Wallis test) and *D. melanogaster* exclusively in yeast media (Figure 2a ; N = 130 total eggs across 11 out of 30 trials, chi-squared = 21.902, p = 1.87e^-5^, Kruskal-Wallis test).

**Figure 2:**
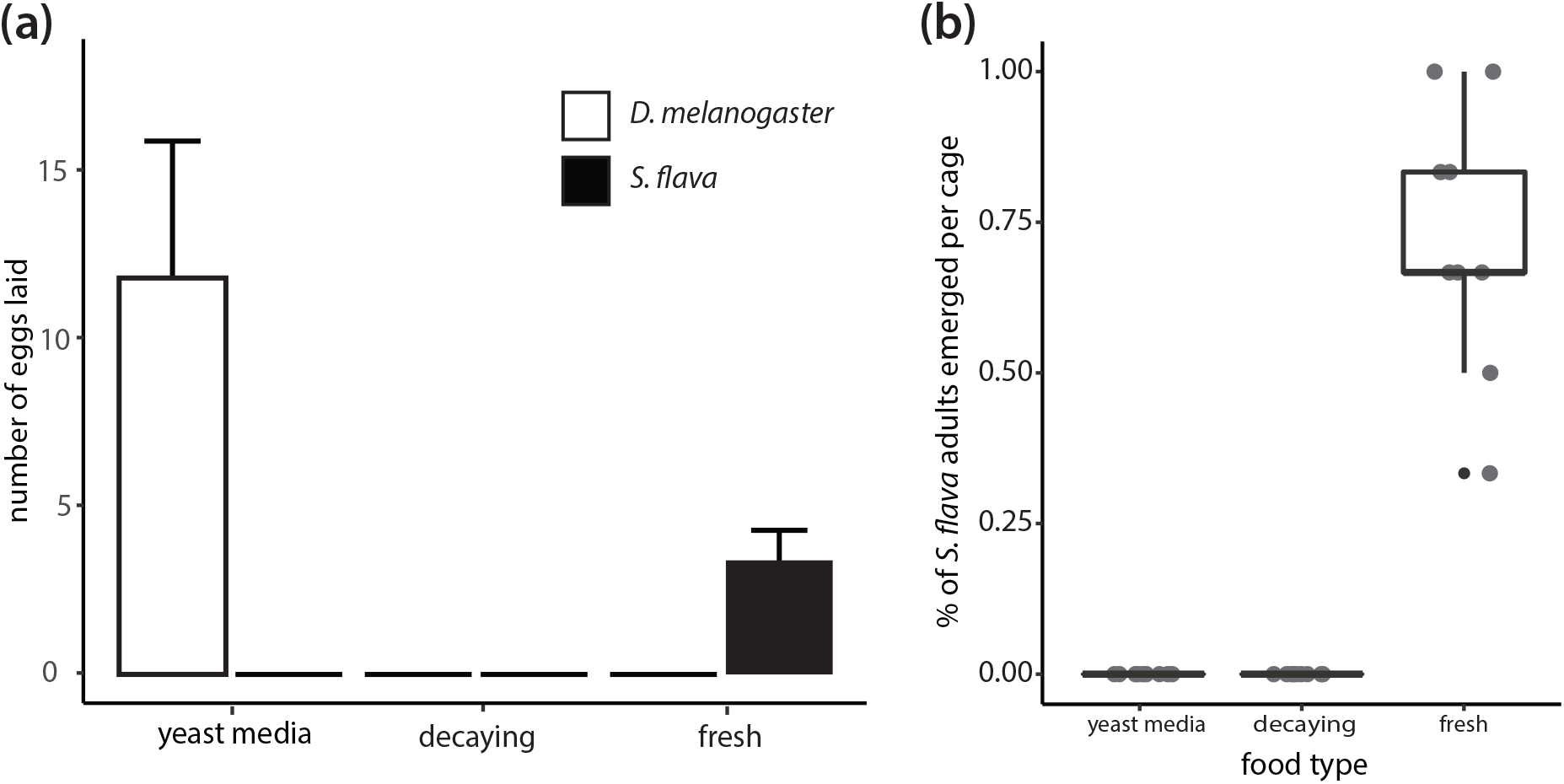
Choice and viability assays confirm that *S. flava* oviposition and development occur on living plants **(a)** Oviposition choice trials. Eggs laid by adult *S. flava* and *D. melanogaster* females in each of three substrates were counted after 24 hours. **(b)** *S. flava* development assay. Per replicate, six second instar *S. flava* were transplanted from fresh plants to individual fresh plants, decaying plants or yeast media tubes. The percentage of adults that emerged is shown. entire leaf and is ready to pupariate. 6. The larva has pupariated in the petiole of the leaf, partially covered by the leaf’s epidermis.

Next, we transferred second instar *S. flava* larvae onto each of the same three substrates. Larvae only completed development to adults in living plant tissue (Figure 2b; N = 9 replicates per substrate, with 6 larvae per replicate, chi-squared = 24.712, P = 9e^-7^, Kruskal-Wallis test).

#### Comparative morphological measurements

We compared gut length divided by body size for four species (*D. melanogaster, S. hsui, S. pallida, S. flava*). We rejected the null hypothesis that average gut length was the same across species (Figure 3a; Supplementary Table 1; P = <0.001). Further, after correcting for multiple comparisons with a Tukey’s HSD test, we found *D. melanogaster* had the largest gut/body ratio compared to the three other species, which all were not different from one another. All ANOVA and Tukey’s HSD results can be found in Supplementary table 1. Next, we compared cephaloskeleton length divided by body size (Figure 3b). We found no differences between *D. melanogaster* and *S. hsui* or *S. pallida* and *S. flava*, but differences between these two species pairings, with *S. pallida* and *S. flava* having relatively larger cephaloskeletons than *D. melanogaster* and *S. hsui* (Supplementary Table 1; P = <0.001). Figures 2d-f show comparisons of the anterior most section of the mouth hook. First, we compared the ratio of mouth hook height to mouth hook length (Figure 3d). We found no differences between *D. melanogaster* and *S. pallida* or *S. hsui* and *S. flava*, but *S. hsui* and *S. flava* had mouth hooks with a longer length from top to bottom compared to *D. melanogaster* and *S. pallida* (Supplementary Table 1; P = <0.001).

**Figure 3:**
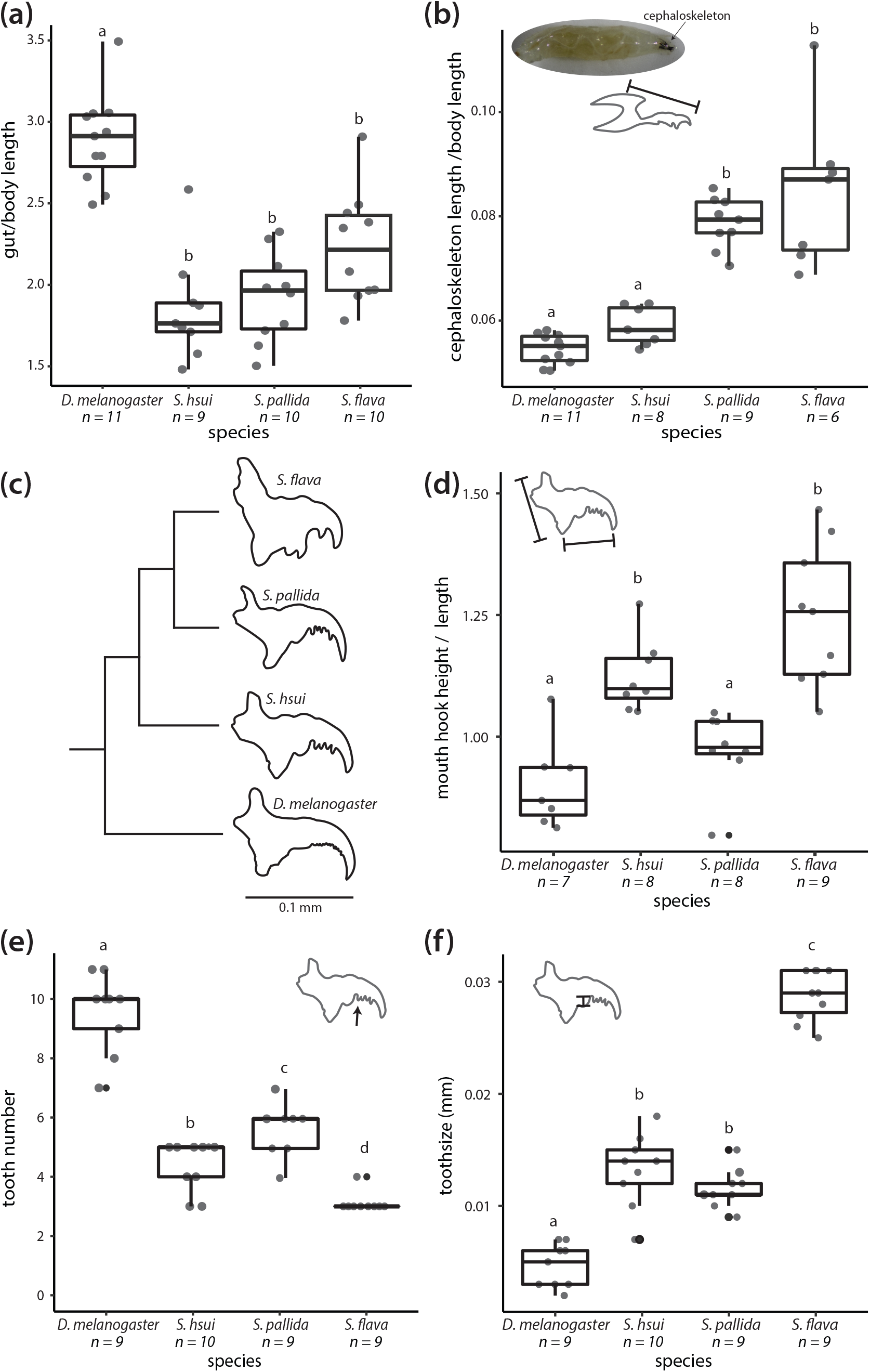
Morphological measurements of characters often associated with herbivory. Data points are offset on the x-axis to aid in visualization. Letters above bars indicate significant differences **(a)** Length of gut relative to body length. **(b)** Length of cephaloskeleton relative to body length; inset photo of a Drosophila melanogaster larvae with cephaloskeleton labeled. **(c)** Representative depiction of mean tooth number, tooth size and phylogenetic relationships of species measured (Peláez et al. 2022). **(d)** Ratio of mouth hook height (here defined as the ventral process of the mouth hook (VPMH) to the dorsal process of the mouth hook) to the length (here defined as the VPMH to the oral process of the mouth hook). **(e)** Mouth hook tooth number. **(f)** Length of the longest tooth.

Finally, we quantified the tooth number and tooth size. Generally, tooth size had an inverse relationship to tooth number, where species with many teeth also had smaller teeth and species with fewer teeth had larger teeth. On average, *D. melanogaster* has the largest number of teeth, followed by *S. pallida, S. hsui*, and *S. flava* (Figure 3e; Supplementary Table 1; P = <0.001). Across the four species measures, *S. pallida* and *S. hsui* have similarly sized teeth that fall between between *D. melanogaster*’s smaller and *S. flava’s* larger teeth (Figure 3f; Supplementary Table 1; P < <0.001). Generally, *S. flava* had fewer, larger teeth, *D. melanogaster* has many small teeth, and *S. hsui* and *S. pallida* fell in the intermediate space for both traits.

#### Sterol Profiling

We measured the composition of sterols in laboratory food sources (Figure 4a) and flies that represent the feeding niches of *S. flava* and its close, non-herbivorous relatives, *D. melanogaster* and *S. hsui* to determine whether there were differences in sterol processing between these species (Figure 4b). To better understand differences between these profiles, we calculated the Bray-Curtis dissimilarity (BCD) of both the food and fly sterol profiles (Figure 4c). Plain, decaying spinach was the most dissimilar from the other three food types, with scores above 0.9, and unsurprisingly, decaying spinach + yeast and plain yeast media were the most similar (Figure 4a, 4c; BCD = 0.26). For fly sterol profiles, *S. hsui* and *D. melanogaster* were the most similar (BCD = 0.6), and *S. flava* and *D. melanogaster* were the least similar (Figure 4b, 4c; BCD = 0.76). Sitosterol dealkylating to cholesterol was most commonly observed in flies eating plant material (Figure 4d)

**Figure 4:**
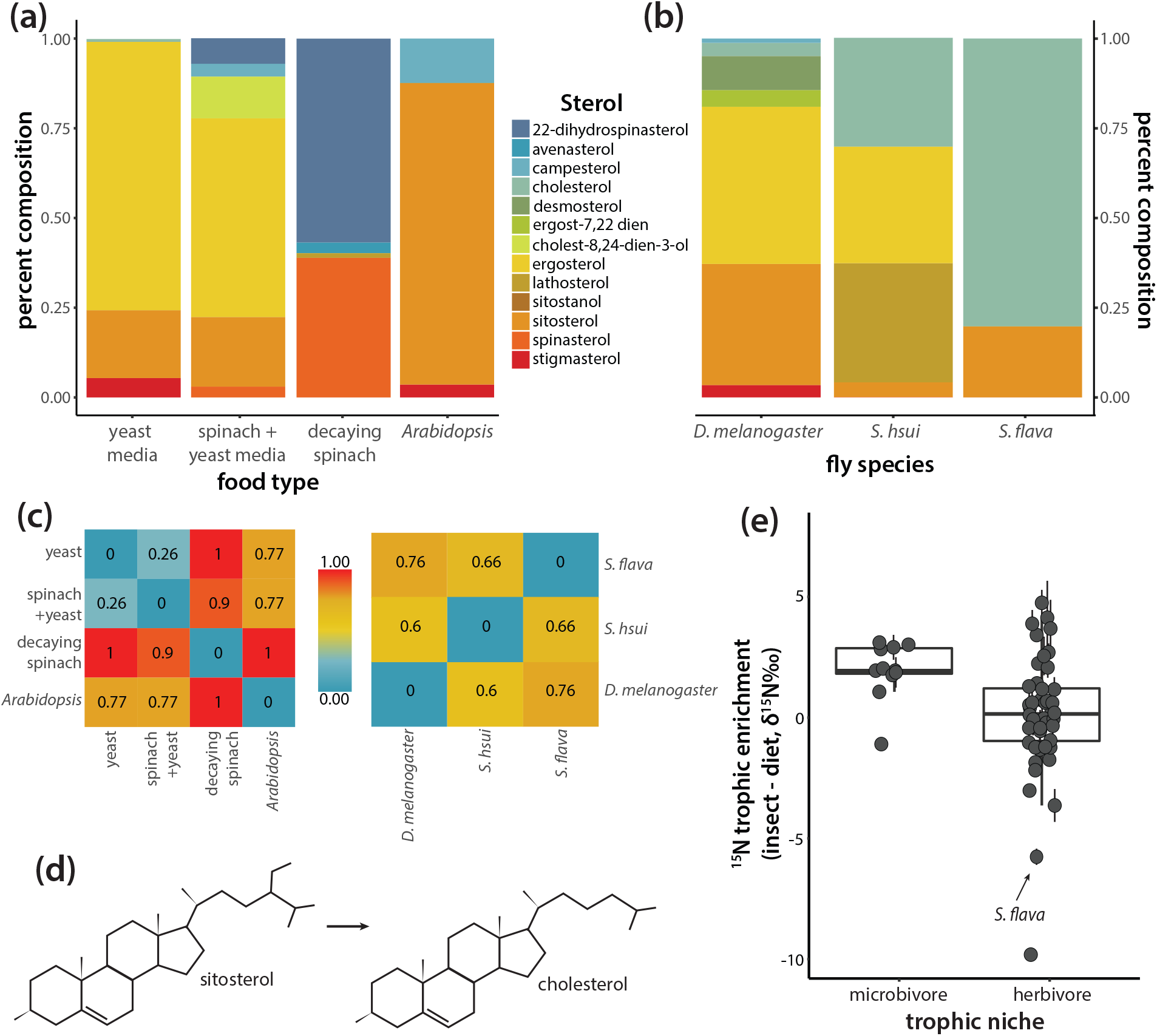
Sterol and nitrogen isotope profiles suggest *S. flava* obtains nutrients from plant leaves **(a)** Sterol composition of laboratory-reared fly food **(b)** Sterol composition of flies reared in the laboratory. *D. melanogaster* eats yeast media, *S. hsui* eats yeast media with decaying spinach, *S. flava* eats fresh *Arabidopsis* plants. **(c)** Bray-Curtis dissimilarity measure of sterol composition of fly food and flies. A 1 is the most dissimilar composition, 0 is completely alike. **(d)** Dealkylation of sitosterol to cholesterol. Sitosterol is the most common type of sterol in plants. **(e)**Trophic enrichment of heavy nitrogen isotopes (δ^15^N, the ratio of ^15^N:^14^N) in insects relative to their feeding substrate, for insects consuming living plant material vs. decaying, microbe-rich plant material. Each circle indicates an averaged observation for a unique combination of insect species and host species or substrate +/- SEM. Data points are offset on the x-axis to aid in visualization. All data except *S. flava* measurements are from the literature. N insects eating decaying plants = 11, N insects eating fresh plants (excluding *S. flava*) = 44; P = .023

#### Isotope profiling

We profiled nitrogen isotope composition in *S. flava* and its *A. thaliana* Col-0 host plants and compared these to additional δ^15^N profiles from the literature to determine whether they were more like an herbivore or non-herbivore δ^15^N trophic enrichment (insect δ^15^N – diet δ^15^N).

Insects feeding on rotting, microbe-rich substrates typically exhibited a two-fold greater trophic enrichment of δ^15^N, than herbivores (Figure 4e; df = 1, Sum sq = 34.3, Mean Sq = 34.34, Sum Sq (Residuals) = 330.4, Mean Sq (residuals) = 6.23, F = 5.51, P = 0.023). Similarly, δ^15^N, was depleted in *S. flava* relative to its host plants, and δ^15^N trophic enrichment was much lower than that of insects feeding on microbe-rich substrates.

## Discussion

The evolution of herbivory is a key innovation in many animal lineages in which it has evolved. Yet, how herbivory evolves is still not well understood. Building on previous work, in this study, we use a natural history approach to investigate the feeding niche of *S. flava*, an herbivorous drosophilid fly that recently evolved from closely related non-herbivorous species. We quantified several key characteristics that aid in understanding trophic niche, including behavior, morphology, and nutritional integration in *S. flava* and three closely related, non-herbivorous species.

The genus *Scaptomyza* is nested within the paraphyletic *Drosophila* clade. *S. flava* attacks the model plant *A. thaliana*, and well annotated reference genomes are available for several *Scaptomyza* species (Kim et al. 2021; Gloss et al. 2019). Although an antagonistic ecological relationship between *Scaptomyza* species and its host plants has been characterized (N. A. Martin 2004; Hackman 1959; Whiteman et al. 2011) the question of its nutritional niche has not been addressed previously. Here, by comparing oviposition choice, morphological measurements, and nutrient integration with closely related non-herbivorous relatives, we found that *S. flava* is an obligate herbivore that consumes and acquires nutrients from living plant tissue.

Since early instar *S. flava* larvae are relatively confined to the location in which they were oviposited, oviposition behavior is key to understanding trophic niche shifts (Bernays and Chapman 1994). In our choice and development trials, *S. flava* showed a strong preference for living plants over decaying plants or yeast media when ovipositing; eggs were laid only on fresh plants (Figure 2a). This is consistent with the observation that adult *S. flava* antennae are responsive to volatiles from leaves but far less so from yeast in olfactory and behavioral assays (Goldman-Huertas et al. 2015; Matsunaga et al. 2022). After transplanting larvae to the three food types (living plants, decaying plants and yeast media), we found a similar trend for larval development to adulthood--adult flies only emerged from living plants (Figure 2b). In contrast, *D. melanogaster* showed a strong preference for yeast media, with eggs laid only on that food type.

Morphological adaptations to food types in multiple life history stages are also necessary for trophic shifts. There have been studies exploring morphological adaptations in the ovipositors of *S. flava* adults that oviposit in leaves (Peláez et al. 2022), and here, we explored morphological adaptations in the herbivorous larval stage of *S. flava*. We used closely related, non-herbivorous groups (*S. pallida, S. hsui*, and *D. melanogaster*) as comparisons. Contrary to the commonly found pattern of longer gut lengths in herbivores compared to non-herbivores, gut length to body ratio was longer in non-herbivorous *D. melanogaster* compared to the other species studied, including the herbivorous *S. flava* (Figure 3a). Cephaloskeleton length was not correlated with herbivory, but we did find morphological differences in mouth hook shape and tooth size between herbivorous and non-herbivorous flies. Herbivorous *S. flava* had fewer, larger teeth on its mouth hooks, which are the mouth organs that mechanically gather food during eating (Figures 3b, 3c, 3d, 3e, 3f). The decrease in tooth number accompanying an increase in tooth size in herbivores is consistent with their utilization of a tougher food source and parallels what was found in carnivorous *D. melanogaster*, which showed a plastic reduction in mouth hook tooth number during shifts from yeast-substrate feeding to carnivory (Vijendravarma, Narasimha, and Kawecki 2013). However, even if putatively herbivorous species show strong oviposition preferences for living plants or morphological shifts consistent with herbivory, it is still possible that these species are obtaining most of their nutrients from microbes living in the leaf. To further understand the source of nutrition for these flies, we measured two metabolic outcomes: sterol profiles and nitrogen stable isotope profiles.

How insects metabolize ingested dietary sterols can provide unique insights to host plant use. For example, the profile of sterols in the bodies of insects is linked to differences in sterol profiles found in insect food, as well as adaptations in dealkylation ability of the species (Janson et al. 2009; Thompson et al. 2013). By comparing the sterol profiles in food to sterol profiles in insects using Bray-Curtis dissimilarity calculations, we obtained a combined picture of these two contributing factors (Figures 4a, 4b, 4c). When comparing herbivores to non-herbivores, we found differences in the efficiency of dealkylation of sitosterol from food sources to cholesterol in insect bodies. Herbivorous *S. flava* had the most dissimilar sterol profile compared to its close non-herbivorous relatives (Figure 4c). *S. flava* was the most efficient at dealkylation of sitosterol, but *S. hsui* is also able to dealkylate this common plant sterol (Figure 4a, 4b, 4d). *S. hsui* eats decaying leaves and is fed a mixture of decaying spinach and yeast media in the lab. The laboratory-fed diet of the two microbe feeding flies, *S. hsui* and *D. melanogaster*, contain the most similar sterol profiles, however the sterol profiles of the insects are still more different than they are alike (Figure 4c). Unsurprisingly, *D. melanogaster* and *S. hsui* both contained ergosterol, which is ubiquitous in fungi, and *S. hsui* can dealkylate the spinach-exclusive sterol spinasterol to lathosterol. Future studies into the mechanistic and evolutionary underpinnings of this are necessary to further interpret the significance of these results as well as to rule out the selective use of cholesterol by these herbivorous larvae (Behmer 2017; Jing and Behmer 2020).

Comparisons of stable isotope composition of nitrogen between insects and putative dietary substrates offer insight into nutrient acquisition: lighter isotopes are preferentially lost through excretion and metabolic activity relative to heavier ones as nutrients move up trophic levels, leading to an enrichment of heavy relative to light nitrogen isotopes (δ^15^N : δ ^14^N) with each trophic step (Perkins et al. 2014). We found δ ^15^N enrichment in insect bodies that reflected the additional trophic step between insect and dietary substrate in microbe feeding insects (i.e., rotting plant tissue → microbe → insect, vs. living plant → insect) (Spence and Rosenheim 2005). Although there is overlap in δ ^15^N enrichment signatures between plant and microbe feeders, increased depletion of δ ^15^N is typical for phloem-feeding herbivorous insects (Wilson, Sternberg, and Hurley 2011; Perkins et al. 2014), and is inconsistent with a diet reliant on plant-associated microbes. Our findings suggest that *S. flava* derives nutrients from the fluid and biomass of living plant tissue (Figure 4e). This is further supported by previous findings that larval feeding and growth rates are not directly enhanced by elevated bacterial loads in their diet, although larvae benefit indirectly from bacterial suppression of plant defenses (Groen et al. 2016).

Overall, the results of our experiments and comparative studies framed by natural history show that *S. flava* is an obligate herbivore which utilizes nutrients from fresh plants to survive. The *S. flava-A. thaliana* interaction system has the potential to facilitate our understanding of how most of the macroscopic biodiversity has evolved and is maintained, as well as fundamental mechanistic questions at the plant-herbivore interface. Future studies are needed to disentangle the adaptive leaps required at the transition to herbivory, as well as the ways herbivorous species diversity is created and maintained once the hurdles to herbivory have been overcome.

## Acknowledgments

The authors would like to thank J.N. Pelaez, R. P. Duncan and M. Karageorgi for assistance with insect care, N.M. Alexandre, C. Miller, and M. Nachman for discussions on morphological measurements and manuscript content, and two anonymous reviewers for constructive feedback on an earlier version of this manuscript. This work was supported by a grant from the NIGMS (R35GM119816) to N.K.W, HHMI Gilliam Fellowship (GT13540) and National Science Foundation Graduate Research Fellowship (DGE 1752814/DGE 2146752) to J.M.A. Any opinion, findings, and conclusions or recommendations expressed in this material are those of the authors(s) and do not necessarily reflect the views of the National Institutes of Health or the National Science Foundation.

