## Supplementary Material for "Insights into the evolution of herbivory from a leaf-mining, drosophilid fly"

### Supplement

*Timing of insect transitions to herbivory*

To estimate the timing of transitions to herbivory in lineages with members that have had their genomes sequenced, we searched for exemplar clades of herbivorous species with reference genomes on InsectBase 2.0 in July of 2022 (Mei et al. 2022). From information available in the literature, categorized them as either obligate herbivore (having at least one life stage where it feeds exclusively on living plant tissues) or facultative herbivore (feeding frequently on plants, but not having obligate herbivory stages). We focused on leaf-feeding insects, and excluded root-feeders, frugivores and granivores. We used stem ages of herbivorous lineages with at least one representative genome assembly from the literature (Wiegmann et al. 2011; Ahrens, Schwarzer, and Vogler 2014; Misof et al. 2014; Wiens, Lapoint, and Whiteman 2015; McKenna et al. 2019). We inferred that the most recent common ancestor of an herbivorous clade was also herbivorous under maximum parsimony and therefore considered these stem ages as estimated dates for origins of herbivory. We visualized the phylogenetic relationships among reference herbivores with iTOL v6 (Letunic and Bork 2021), and included the herbivorous spider mite (*Tetranychus urticae*) as an outgroup (Rota-Stabelli, Daley, and Pisani 2013; Xue et al. 2017).

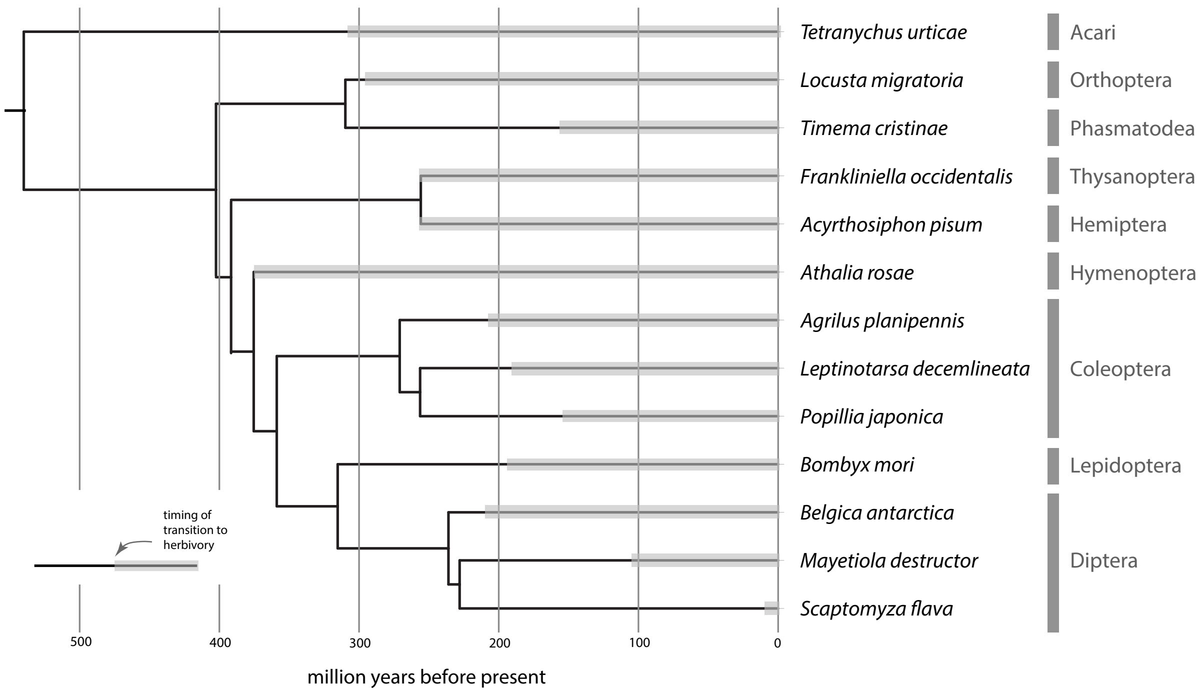


**Supplementary Figure 1:** The timing of transitions to herbivory, inferred from the reported diet niche of related families and orders, in the ancestors of exemplar clades of terrestrial herbivorous arthropods with sequenced genomes. Further details and source data are included in Supplementary Table 1.

Data file: HerbivoryPaper_ OriginOfHerbivory_20220905.csv

UC Berkeley yeast media ingredients:

water, agar, cornmeal, yeast, sucrose, sodium tartrate (4 hydrate), $CaCl_2$ (2 hydrate), molasses, tegosept in 95% EtOH, and propionic acid

Bloomington yeast media ingredients:

water, cornmeal, light corn syrup, malt extract, yeast, soy flour, agar, antimicrobial agents

*Sterol extraction and processing*

We extracted samples using equal volumes (8 milliliters) of chloroform, 70% methanol, and water. Following a 12 hour incubation, the chloroform layer containing the free and base hydrolysable sterols was removed and evaporated to a reduced volume (less than 100 ul). We set aside the 70% methanol:water layer containing glycosylated sterols for additional processing and sterol identification.

We resuspended the reduced volume chloroform layer (less than 100 ul but not dryness) in 5 mls of hexane and 5 mls of 70% methanol/water combined. We removed the original hexane layer following separation of the phases. We added a second 5 ml volume of hexane to the original 70% methanol/water layer and mixed vigorously. After separation, we removed the second hexane layer, combined the two hexane volumes, and evaporated to a reduced volume – less than 100 ul.

To release base hydrolysable sterol esters, we resuspended the hexane fraction in 8 ml of 70% methanol/water containing 5% KOH w/v (with sonication) and incubated at 70°C in a water bath for 1.5 hours with constant shaking (Feng et al. 2015). After base saponification, we added 3 ml of water and 5 ml of water equilibrated hexane, shook vigorously and vortexed for 1 minute. We then left samples for at least 6 hours, until the layers separated, or centrifuged for 2 minutes to assist in phase separation (MSE GT-2 centrifuge equipped with a swinging bucket rotor – at 1,500 rpms - VWR). We removed the hexane layer (containing the released sterols). We added 4 ml of hexane to the remaining methanol/water, repeated the above steps, and again removed the hexane layer. Using the above separation protocols, we combined the two hexane volumes and backwashed to neutrality against an equal volume of water, with neutrality determined by PHydrion Mikro Ion paper (Micro Essential Laboratories, Brooklyn, NY). We concentrated the neutral hexane fraction by evaporating to a minimal volume (less than 100 ul). All of the described procedures used solvents that were pre-equilibrated to the paired solvent. We followed the procedure described in Feng et al., 2015 to release the glycosylated sterols.

| | Adjusted P values | | | | |
|---|---|---|---|---|---|
| | Gut/body | Cephalo-skeleton/ body | Mouth hook height/length | Mouth hook tooth # | Mouth hook toothsize |
| Df | 3 | 3 | 3 | 3 | 3 |
| Df, residuals | 36 | 31 | 28 | 33 | 33 |
| Sum Sq | 6.946 | 0.006 | 0.584 | 0.003 | 217.80 |
| Sum Sq, residuals | 3.358 | 0.002 | 0.299 | 1.85e-4 | 27.12 |
| Mean Sq | 2.315 | 0.002 | 0.195 | 1.0e-03 | 72.6 |
| Mean Sq, residuals | 0.0933 | 5.46e-5 | 0.011 | 5.6e-06 | 0.82 |
| F-value | 24.82 | 34.7 | 18.23 | 178 | 88.33 |
| Pr(>F) | <0.001 | <0.001 | <0.001 | <0.001 | <0.001 |
| | **Tukey's HSD species comparisons** | | | | |
| *D. melanogaster / S. hsui* | <0.001 | 0.580 | 0.001 | <0.001 | <0.001 |
| *D. melanogaster / S. pallida* | <0.001 | <0.001 | 0.544 | <0.001 | <0.001 |
| *D. melanogaster / S.flava* | <0.001 | <0.001 | <0.001 | <0.001 | <0.001 |

| | | | | | |
|---|---|---|---|---|---|
| *S. pallida* / *S. hsui* | 0.955 | <0.001 | 0.033 | 0.019 | 0.454 |
| *S. hsui* / *S. flava* | 0.051 | <0.001 | 0.086 | 0.028 | <0.001 |
| *S. pallida* / *S. flava* | 0.134 | 0.367 | <0.001 | <0.001 | <0.001 |

**Supplementary Table 2.** P values of morphological measurements, adjusted for multiple comparisons

Ahrens, Dirk, Julia Schwarzer, and Alfried P. Vogler. 2014. “The Evolution of Scarab Beetles Tracks the Sequential Rise of Angiosperms and Mammals.” *Proceedings. Biological Sciences / The Royal Society* 281 (1791): 20141470.

Feng, Simin, Songbai Liu, Zisheng Luo, and Kaichen Tang. 2015. “Direct Saponification Preparation and Analysis of Free and Conjugated Phytosterols in Sugarcane (Saccharum Officinarum L.) by Reversed-Phase High-Performance Liquid Chromatography.” *Food Chemistry* 181 (August): 9–14.

Letunic, Ivica, and Peer Bork. 2021. “Interactive Tree Of Life (ITOL) v5: An Online Tool for Phylogenetic Tree Display and Annotation.” *Nucleic Acids Research* 49 (W1): W293–96.

McKenna, Duane D., Seunggwan Shin, Dirk Ahrens, Michael Balke, Cristian Beza-Beza, Dave J. Clarke, Alexander Donath, et al. 2019. “The Evolution and Genomic Basis of Beetle Diversity.” *Proceedings of the National Academy of Sciences of the United States of America* 116 (49): 24729–37.

Mei, Yang, Dong Jing, Shenyang Tang, Xi Chen, Hao Chen, Haonan Duanmu, Yuyang Cong, et al. 2022. “InsectBase 2.0: A Comprehensive Gene Resource for Insects.” *Nucleic Acids Research* 50 (D1): D1040–45.

Misof, Bernhard, Shanlin Liu, Karen Meusemann, Ralph S. Peters, Alexander Donath, Christoph Mayer, Paul B. Frandsen, et al. 2014. “Phylogenomics Resolves the Timing and Pattern of Insect Evolution.” *Science* 346 (6210): 763–67.

Rota-Stabelli, Omar, Allison C. Daley, and Davide Pisani. 2013. “Molecular Timetrees Reveal a Cambrian Colonization of Land and a New Scenario for Ecdysozoan Evolution.” *Current Biology: CB* 23 (5): 392–98.

Wiegmann, Brian M., Michelle D. Trautwein, Isaac S. Winkler, Norman B. Barr, Jung-Wook Kim, Christine Lambkin, Matthew A. Bertone, et al. 2011. “Episodic Radiations in the Fly Tree of Life.” *Proceedings of the National Academy of Sciences of the United States of America* 108 (14): 5690–95.

Wiens, John J., Richard T. Lapoint, and Noah K. Whiteman. 2015. “Herbivory Increases Diversification across Insect Clades.” *Nature Communications* 6 (September): 8370.

Xue, Xiao-Feng, Yan Dong, Wei Deng, Xiao-Yue Hong, and Renfu Shao. 2017. “The Phylogenetic Position of Eriophyoid Mites (Superfamily Eriophyoidea) in Acariformes Inferred from the Sequences of Mitochondrial Genomes and Nuclear Small Subunit (18S) RRNA Gene.” *Molecular Phylogenetics and Evolution* 109: 271–82.
